# Cadmium toxicity to the human gut microbiome varies depending on composition

**DOI:** 10.1101/2025.11.19.689308

**Authors:** Carmen E. Perez-Donado, Sujun Liu, Javier Seravalli, Jennifer M. Auchtung, Devin J. Rose

## Abstract

Cadmium (Cd(II), hereafter Cd) is a toxic heavy metal with detrimental impacts on the gut microbiota. We investigated the effects of acute Cd exposure on human fecal microbiotas using 24-hour *in vitro* cultures from 21 healthy adult donors. Regression analysis of butyrate production in the absence (-Cd) versus presence (+Cd) of Cd identified three categories of microbial responses: sensitive, intermediate, and resilient. Under Cd stress, sensitive microbiomes exhibited significant decreases in butyrate coupled with elevated acetate and lactate production, while resilient microbiomes did not show significant changes in butyrate and exhibited attenuated increases in lactate compared with sensitive microbiomes. Several genera were significantly different between sensitive and resilient communities after exposure to Cd, but the most striking was *Anaerostipes*. Network analysis revealed a significantly greater disruption of microbial interactions in sensitive communities compared with resilient. In resilient communities, butyrate production was primarily associated with *Faecalibacterium* in the absence of Cd and *Anaerostipes* in the presence of Cd. Furthermore, supplementation of sensitive microbiota with *Anaerostipes* species restored butyrate production in the presence of Cd. These findings highlight distinct gut microbial responses to acute Cd exposure and provide a foundation to investigate microbiota features underlying Cd sensitivity or resilience.

**IMPORTANCE:** Cadmium is a widespread environmental contaminant that reaches the human intestine, where it can disrupt the gut microbial community and negatively impact digestive and systemic health. However, this study demonstrates that human gut microbiomes vary in their responses to cadmium exposure: sensitive communities exhibit losses of beneficial organisms, particularly butyrate-producing taxa that contribute to intestinal integrity and metabolic balance, whereas resilient communities retain microorganisms with this key functional capacity. *Anaerostipes* appeared to be involved, at least in part, with Cd resilience. This work advances our understanding of how gut microbial functions may mitigate the adverse effects of cadmium exposure by identifying the compositional features that distinguish sensitive from resilient microbiomes. These findings highlight the importance of elucidating microbiome-mediated mechanisms that help sustain host health and lay the groundwork for deeper mechanistic studies aimed at mitigating cadmium toxicity.

## Introduction

Cadmium (Cd(II), hereafter Cd), a toxic heavy metal prevalent in various environmental sources, poses significant health risks primarily through dietary intake (1). Despite its relatively low bioavailability via the gastrointestinal tract (3-8%)(2), cadmium’s long biological half-life causes accumulation in the renal cortex with minimal excretion that can have profound impacts on human health, and contribute to a spectrum of disorders affecting multiple organs, including the skeleton, kidneys, gastrointestinal tract and metabolism (3–5) (6). Initially sequestered by metallothioneins (MTs), Cd is released upon MT degradation or saturation, triggering oxidative stress and tissue damage (7, 8). A systematic review linked chronic oral Cd exposure to increased risks of cancers (endometrial, melanoma, breast, prostate, bladder, gastric, and pancreatic) and bone disorders, including osteoporosis (9).

Because the bioavailability of Cd is low, the majority of ingested Cd survives upper gastrointestinal transit and can interact with the gut microbiota. Recent research highlights the potential of Cd to alter microbial metabolism, compromise intestinal barrier function, and enhance Cd absorption, thereby exacerbating metabolic complications (10–16). Oral Cd intake has been associated with decreased abundance of key beneficial microbes such as *Lactobacillus, Bifidobacterium*, *Prevotella,* and *Lachnoclostridium*. A decline in butyrate-producing taxa, such as *Blautia* and *Clostridium_XlVa*, has also been reported, accompanied by an enrichment of opportunistic pathogens such as *Escherichia coli*/*Shigella* (12, 14, 17). These shifts in microbial composition, along with elevated levels of lipopolysaccharide (LPS), have been implicated in liver, kidney, and reproductive dysfunction, suggesting broader effects beyond the gut. The consistent depletion of short-chain fatty acid (SCFA)-producing bacteria and the resulting increase in gut permeability and inflammation further reinforce the link between Cd-induced dysbiosis and host metabolic disturbances (11). Notably, most of these findings stem from studies conducted in animal models, primarily rodents, which have key differences in gut microbiota composition from humans, highlighting the need to evaluate whether similar effects occur in the human gut microbiome (18–22).

Given these observations, understanding the impact of Cd on SCFA production is particularly critical, as SCFAs are essential microbial metabolites that help maintain gut health. Produced mainly through the fermentation of dietary carbohydrates, SCFAs play vital roles in regulating gene expression, modulating immune responses, and influencing host metabolism (23, 24). Cd exposure has been shown to impair microbial carbohydrate metabolism pathways that are critical for SCFA synthesis (11). Moreover, Cd-induced dysbiosis is marked by significantly reduced microbial richness and the loss of key SCFA-producing taxa, particularly butyrate producers. These disruptions ultimately lead to lower SCFA levels, which may contribute to both intestinal and systemic dysfunction (11, 14).

In a recent study, we investigated the effects of Cd exposure on human gut microbiota using minibioreactor arrays (MBRAs), a continuous flow model of the colon (25). Our findings revealed distinct responses to Cd among microbiomes from two healthy individuals, highlighting variation in microbial composition, function, and SCFA production, particularly butyrate, and suggesting differing levels of Cd sensitivity. Notably, butyrate-producing bacteria appeared especially susceptible to Cd in the sensitive community. These findings suggested that human gut microbiotas may differ in their phenotypic response to Cd exposure. However, these studies also demonstrated that further research was needed to determine whether consistent Cd-sensitive and Cd-resilient microbiomes could be identified across a broader population and to elucidate whether butyrate production was affected in other sensitive microbiomes under Cd stress. Building upon this foundation, the objective of this study was to determine the proportion of human gut microbiomes exhibiting Cd sensitivity or resilience and identify conserved compositional and functional changes between resilient and sensitive communities. We hypothesized that variations in gut microbiota composition modulate the community’s susceptibility or resilience to Cd exposure in terms of SCFA production, carbohydrate degradation, Cd binding capacity, and the structure of microbial interaction networks. We observed clear distinctions between Cd-sensitive and Cd-resilient human gut microbiomes, with resilient microbiomes continuing to produce butyrate under Cd-stress, which correlated with elevated levels of the butyrate-producing genus, *Anaerostipes* and predicted enrichment of metabolic pathways for stress resistance. In contrast, butyrate metabolism was significantly impaired in Cd-sensitive microbiomes, which lacked butyrate-producing *Anaerostipes* species during Cd-stress and exhibited more significant changes in microbial interaction networks than those observed in Cd-resilient microbiomes. Further supporting the central role of *Anaerostipes* in maintaining butyrate production during Cd-stress, we observed that supplementation of sensitive microbiota with *Anaerostipes* species restored butyrate production in the presence of Cd. Altogether, these studies provide new insights into potential mechanisms underlying microbiota-specific differences in Cd-sensitivity.

## Results

### Cd alters microbial metabolite production

To evaluate the effects of Cd on human gut microbial communities, we analyzed SCFA, lactate, and branched-chain fatty acid (BCFA) production of the *in vitro* cultures before and after fermentation in Cd-free (-Cd) and Cd-containing (+Cd) media (Figure 1). Cd had effects on butyrate and lactate concentrations across the largest number of microbiomes, although responses of individual microbiomes to Cd varied. The inverse effects of Cd exposure on butyrate and lactate concentrations across microbiomes was consistent with previous observations that lactate is often a transient, intermediate metabolite of gut microbiota fermentation of carbohydrates to butyrate.

**Figure 1.**
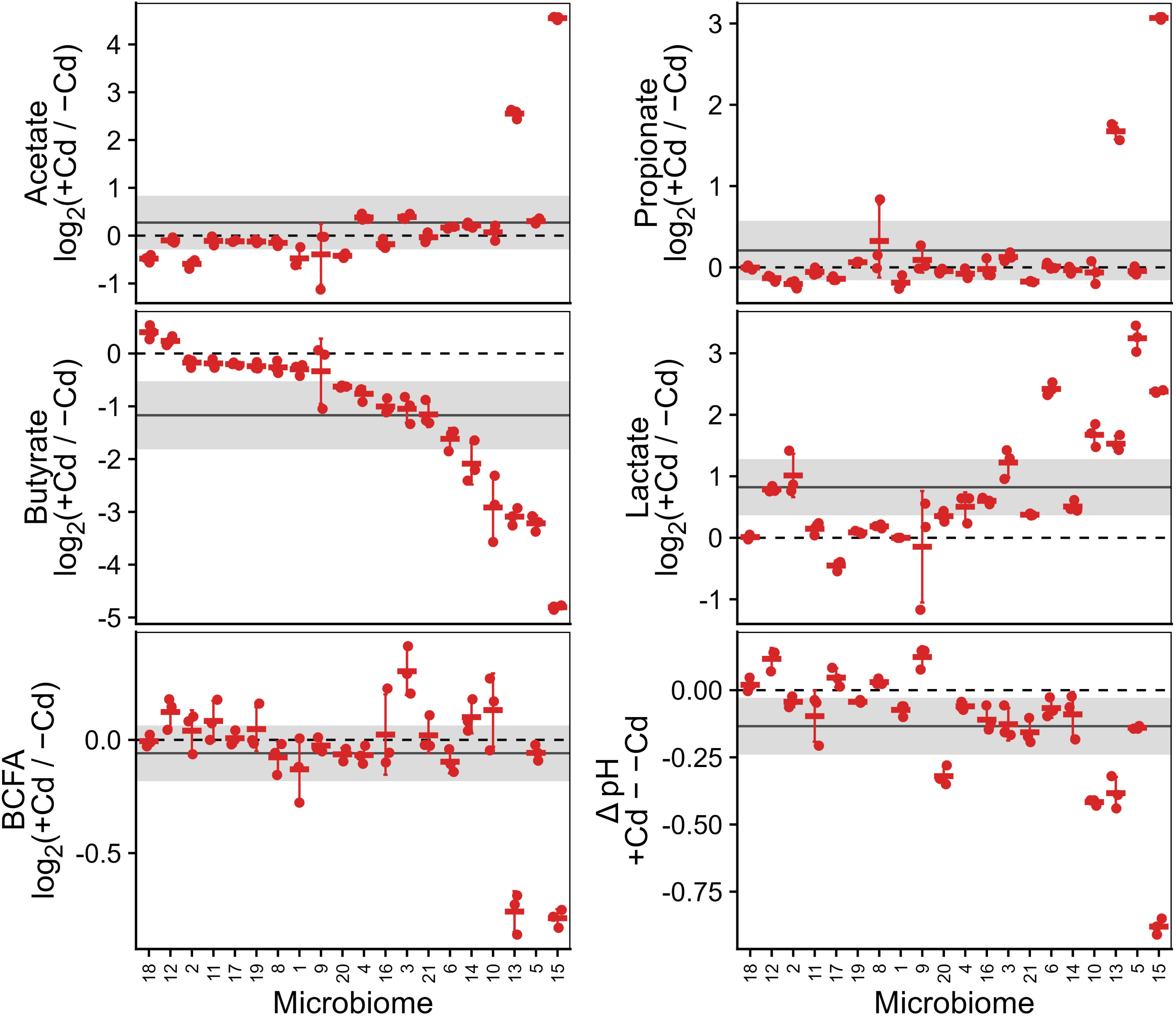
Cd exposure generally reduces butyrate and increases lactate concentrations with substantial variability among microbiomes. Log_2_-fold change in short chain fatty acids (SCFA), branched chain fatty acids (BCFA), and lactate concentrations and change in pH were calculated by comparing cadmium-treated (+Cd) and untreated (-Cd) *in vitro* cultures from 20 human gut microbiomes (n = 3 technical replicates per microbiome). Each microbiome is indicated by a unique number and is ordered by descending log2-fold change in butyrate. The solid horizontal line indicates the mean log_2_-fold change per metabolite across all microbiomes, with the shaded region representing the 95% confidence interval. The dotted line marks zero on the y-axis (no change). Crossbars and error bars denote the mean ± standard deviation for each microbiome.

### Changes in butyrate production in the presence of Cd separate microbiomes into sensitive and resilient groups

Given the large number of microbiomes that exhibited decreased butyrate production during Cd-stress, we classified microbial communities based on butyrate production. As the log_2_-fold change analysis only quantified change in butyrate production and did not differentiate microbiomes with high butyrate production from those with low butyrate production, we conducted a linear regression analysis of butyrate produced with and without Cd. This analysis showed a linear relationship between butyrate production in the presence and absence of Cd, indicating a consistent trend of reduced butyrate production across most of the microbiomes tested (Figure 2A). Among the 20 microbiomes analyzed, six microbiomes were positioned above the regression line, outside the 95% confidence interval. These microbiomes were categorized as ‘resilient’ because they had higher butyrate production under Cd stress than would be predicted based on the regression across all microbiomes. Conversely, five microbiomes, situated below the linear regression line, were classified as ‘sensitive,’ as they exhibited lower butyrate production under Cd stress than would be predicted based on the data across all microbiomes. Importantly, these classifications are operationally defined based on microbiome responses to the single Cd concentration tested in this study, and therefore may vary under different Cd exposure levels.

**Figure 2.**
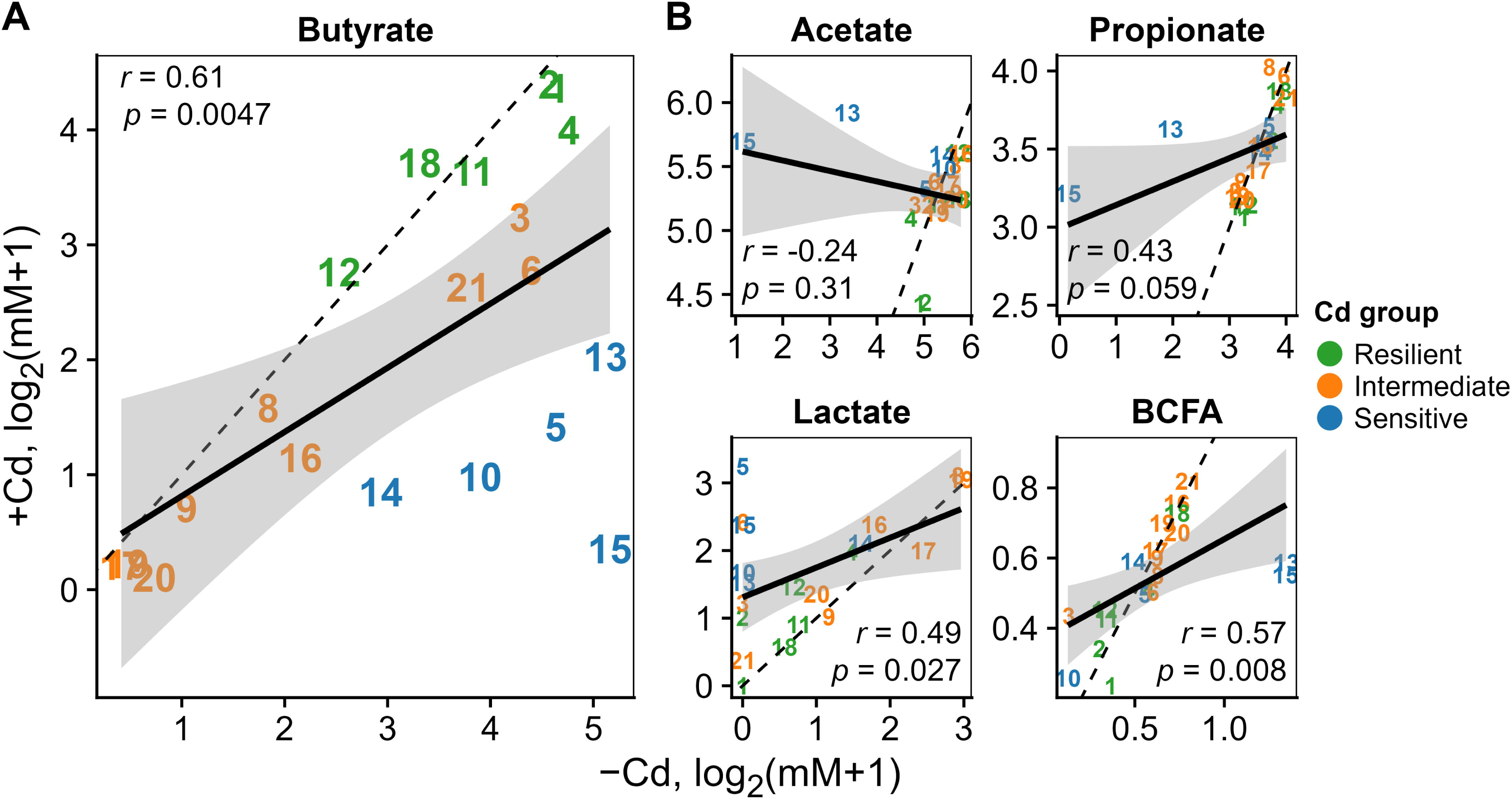
Correlation analysis between butyrate production with (+Cd) and without (-Cd) cadmium reveals Cd-sensitive and Cd-resilient microbiomes. Butyrate (A) and other metabolite (B) concentrations measured after 24 h with (+Cd) and without (-Cd) cadmium. The mean (n=3) for each microbiome is labeled with the microbiome number, colored by Cd-sensitivity group based on butyrate production. The black solid line and shaded region show the regression line and 95% confidence interval of the regression line; the dotted line is y = x.

We also analyzed the correlation between the production of other metabolites in the presence and absence of Cd (Figure 2B). For acetate, propionate, and BCFA, most microbiomes exhibited no difference between concentrations measured after culturing in the presence versus absence of Cd (i.e., fell on the dotted line corresponding to Y=X); the few outliers observed belonged to the sensitive group. For lactate, there were similarly no differences in concentrations measured after culturing in the presence versus absence microbiomes in the resilient and intermediate microbiomes. However, there were significant differences in most of the sensitive microbiomes as all but one microbiome in the Cd-sensitive group except one had very low measurable lactate in the absence of Cd, but had among the highest lactate concentrations in the presence of Cd.

### Cd sensitivity groups differed in metabolite production and carbohydrate utilization in the presence of Cd

We then analyzed differences in metabolite production and carbohydrate utilization across Cd-sensitive and Cd-resilient microbiomes with or without Cd-stress (Figure 3). In the absence of Cd stress, no significant differences were found between Cd-sensitive and Cd-resilient microbiomes, although we observed higher levels of inter-individual variation in acetate, propionate, and BCFA production within the Cd-sensitive microbiomes in the absence of Cd, primarily driven by microbiomes 13 and 15. In contrast, under Cd stress, we observed that the significant decreases in butyrate concentration in the sensitive microbiotas were accompanied by significant increases in acetate and lactate concentrations, which were also significantly higher than those observed in Cd-resilient communities in the presence of Cd.

**Figure 3.**
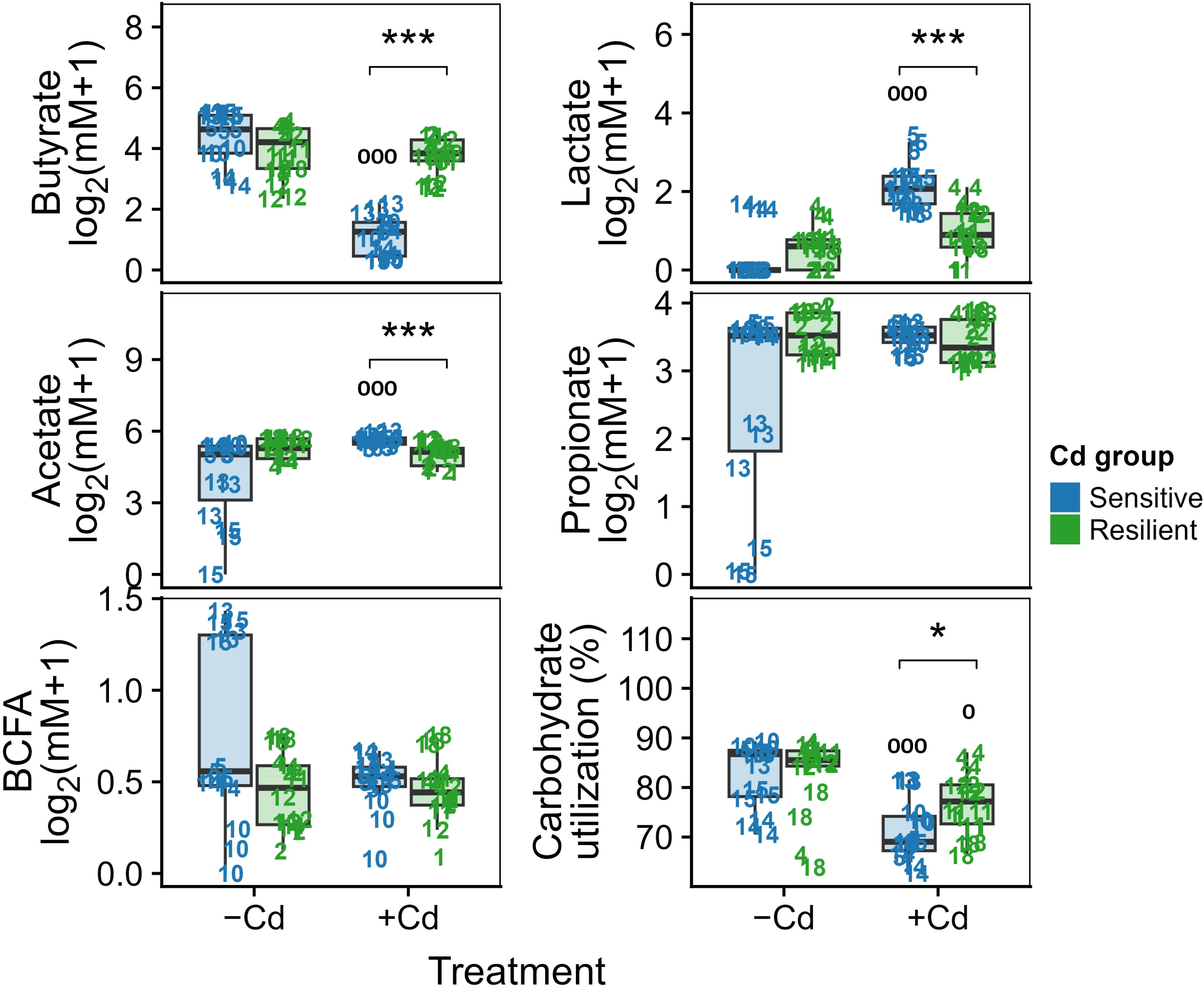
Analysis of differences in metabolite production and carbohydrate utilization between Cd-sensitive and Cd-resilient groups reveals significant changes in Cd-sensitive groups in the presence of Cd. Metabolite and carbohydrate utilization measured after 24 h with (+Cd) and without (-Cd) cadmium was plotted for Cd-sensitive and Cd- resilient communities. Asterisks denote significant differences between sensitive and tolerant microbiomes (*, p<0.05); “0” annotations indicate significant differences from -Cd within each group (^00^, p<0.01; ^000^, p<0.001) as determined by Wilcoxon test with Holm’s p-value adjustment. Numerical data for each microbiome can be found in Supplementary File 1.

For carbohydrate degradation, both Cd-sensitive and Cd-resilient microbiomes exhibited decreased carbohydrate utilization in the presence of Cd versus its absence (Figure 3). However, carbohydrate utilization decreased to a greater extent in sensitive communities and was significantly lower than that observed in resilient microbiotas in the presence of Cd.

### Sensitive and resilient microbiomes had small differences in overall microbiota composition

Fecal microbiota composition was characterized in resilient and sensitive microbiomes at baseline prior to cultivation. Analysis of weighted and unweighted UniFrac dissimilarities revealed small, but significant, differences in overall microbiota composition and structure (Figure 4A), with significant overlap between sensitive and resilient microbiota on principal coordinates biplots. There were no significant differences in species richness (observed ASVs) or microbial diversity (Shannon’s index) between resilient and sensitive communities (Figure 4B). Sensitive microbiotas had higher abundances of members of the Actinobacteria (Actinomycetota) phylum, primarily driven by higher levels of *Bifidobacterium* and *Collinsella* species (Figure 4C; Supplementary Table). Resilient microbiotas contained higher abundances of the Deltaproteobacteria class, specifically from *Bilophila*. Additionally, four genera from the Firmicutes (Bacillota) phylum were present in significantly higher abundances in resilient fecal microbiotas: *Erysipelotrichaceae* UCG-003, *Ruminiclostridium* 9, *Ruminococcaceae* UCG-002, and *Phascolarctobacterium*.

**Figure 4.**
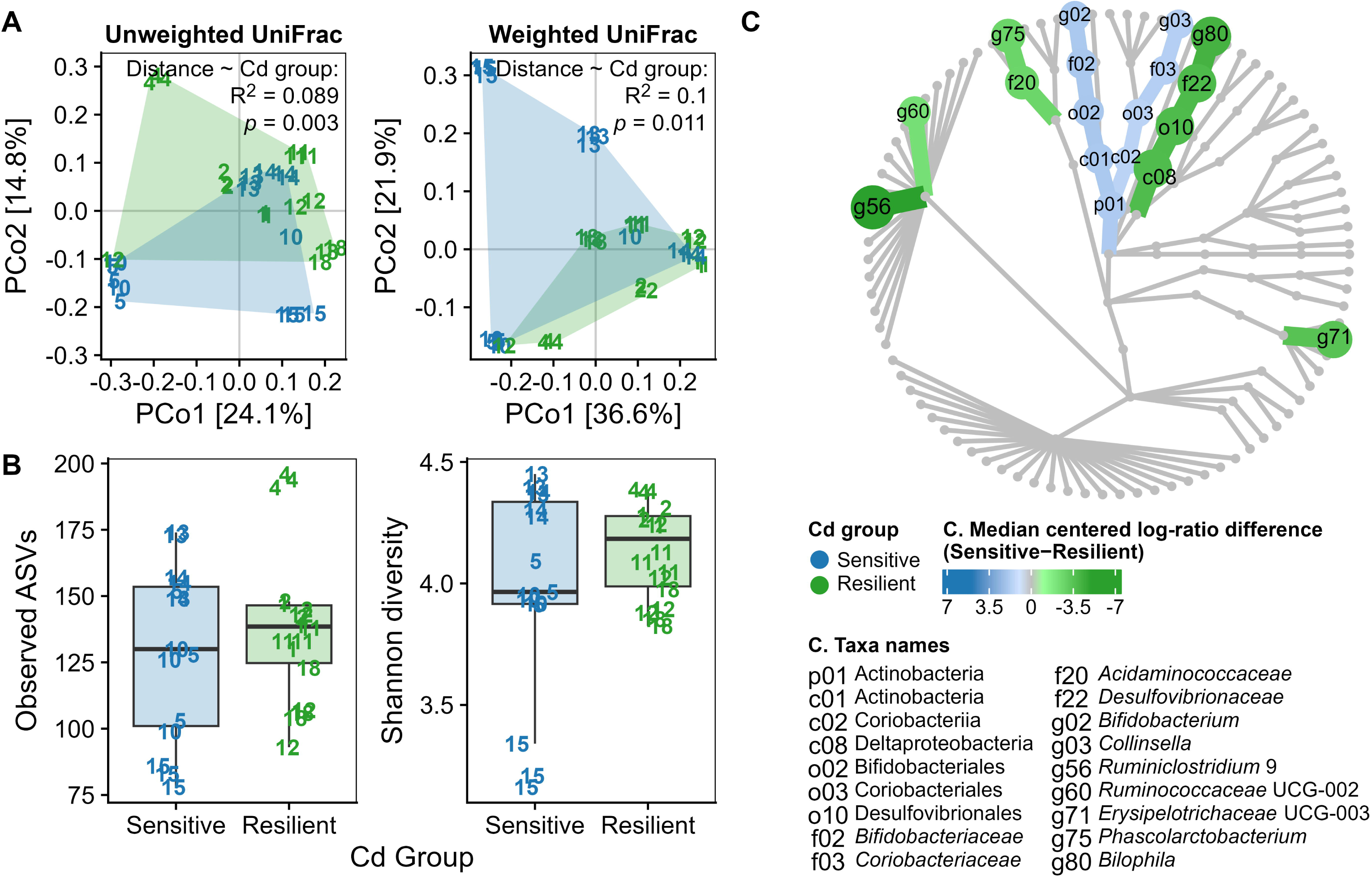
Characterization of fecal microbiotas prior to cadmium exposure. A) Principal coordinates biplots based on unweighted and weighted UniFrac dissimilarities with inset statistics showing the proportion of variance explained by Cd group (resilient or sensitive, defined post hoc). B) Microbiota richness and diversity of sensitive and resilient fecal microbiotas. C) Heat tree showing taxa with significant differences between Cd-sensitive and Cd-resilient microbiotas. Taxa are colored by the Cd group for which they were significantly enriched, with the darkness of the color and size of the node indicating the effect size and gray nodes indicating no significant differences. Wilcoxon’s test with Benjamini-Hochberg p-value adjustment implemented after centered log-ratio transformation of raw read counts implemented within the ALDEx2 R-package.

### Cd had differential effects on microbiota composition in sensitive and resilient microbiotas

We investigated whether microbial diversity and community composition differed after cultivation with or without Cd stress in sensitive or resilient communities. Similar to the baseline fecal communities, permutational multivariate analysis of variance using distance matrices (PERMANOVA) showed that Cd group accounted for only a small proportion of the variance in composition among communities in the presence or absence of Cd (unweighted UniFrac: -Cd: R^2^ = 0.09, p=0.001; +Cd: R^2^ = 0.07, p=0.02; weighted UniFrac: -Cd: R^2^ = 0.039, p=0.34; +Cd: R^2^ = 0.12, p=0.001). We also observed no differences between Cd-sensitive and Cd-resilient microbiomes for weighted or unweighted UniFrac dissimilarities with or without Cd (Figure 5A). Likewise, no differences in species richness or Shannon diversity were observed between Cd-sensitive and Cd-resilient groups after 24 h of culturing without or with Cd (Figure 5B).

**Figure 5.**
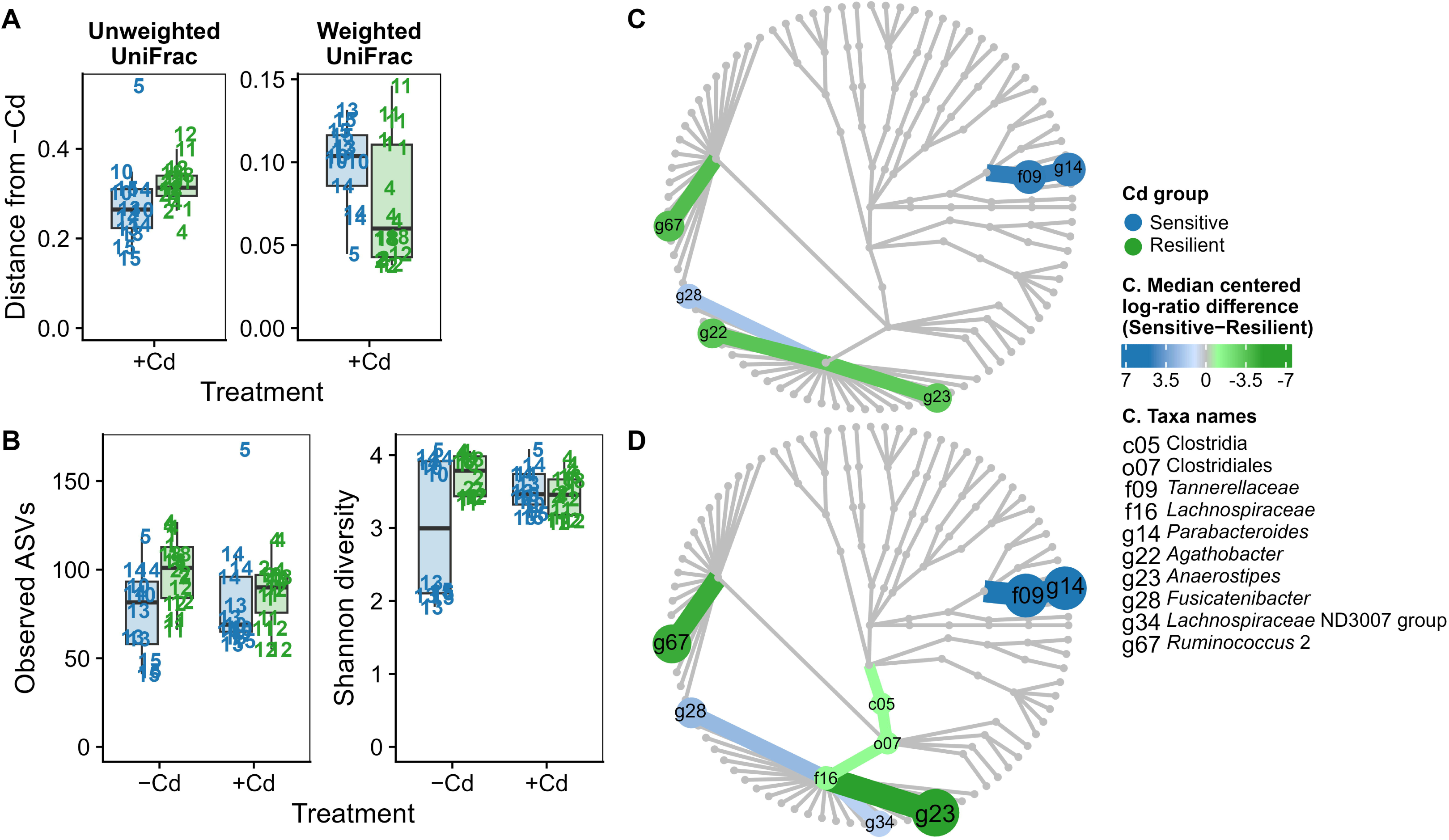
Differences in microbiota composition between sensitive and resilient microbiotas observed after fermentation with (+Cd) and without (-Cd) cadmium. A) Unweighted and Weighted UniFrac dissimilarity between +Cd and -Cd conditions for each *in vitro* culture after 24 h of fermentation for Cd-sensitive and Cd-resilient microbiomes. B) Microbial richness and Shannon diversity after 24 h of fermentation with (+Cd) or without (Cd-) for Cd-sensitive and Cd-resilient microbiomes. C, D) Heat trees showing taxa with significant differences between Cd-sensitive and Cd-resilient microbiotas without Cd (C) or with Cd (D). Heat trees are colored by the Cd group for which they are significantly enriched, with the darkness of the color and size of the node indicating the effect size and gray nodes indicating no significant differences. No statistical differences were found in panels A and B as determined by Wilcoxon’s test with Holm’s p-value adjustment. Statistical significance of panels C and D were determined by Wilcoxon’s test with Benjamini-Hochberg p-value adjustment implemented after centered log-ratio transformation of raw read counts implemented within the ALDEx2 R-package.

After 24 h of culture in the absence of Cd, we observed Cd-sensitive communities had higher abundance of *Parabacteroides* and one genus in the *Lachnospiraceae* family: *Fusicatenibacter* (Figure 5C; Supplementary Table). In contrast, the Cd-resilient communities contained higher abundances of *Ruminococcus* 2 as well as two *Lachnospiraceae* genera: *Agathobacter* and *Anaerostipes*.

Significant differences between Cd-sensitive and Cd-resilient communities observed without Cd were also seen with Cd, but the differences were generally larger (Figure 5D). We also observed significantly increased levels of the *Lachnospiraceae* family in Cd-resilient communities, largely due to greater differences in levels of *Anaerostipes* between resilient and sensitive communities.

To further investigate taxonomic differences between Cd-sensitive and Cd-resilient communities, we identified all genera that differed significantly between microbiome groups with or without Cd. We plotted the abundance of these six genera at baseline and after 24 h of culture with or without Cd (Figure 6A). Two genera, *Anaerostipes* and *Ruminococcus* 2, were significantly higher in Cd-resilient communities in both the presence and absence of Cd. Two genera, *Parabacteroides* and *Fusicatenibacter*, were significantly lower in Cd-resilient communities without and with Cd. Two genera also increased significantly compared with baseline under Cd stress: *Parabacteroides* in the sensitive communities and *Anaerostipes* in the resilient communities. These analyses provided further support for our earlier observation that *Anaerostipes* levels were significantly higher in Cd-resilient communities in culture, and further showed that *Anaerostipes* levels increased significantly from baseline during culture of Cd-resilient communities, whereas no significant changes in *Anaerostipes* levels were observed from baseline in Cd-sensitive communities.

**Figure 6.**
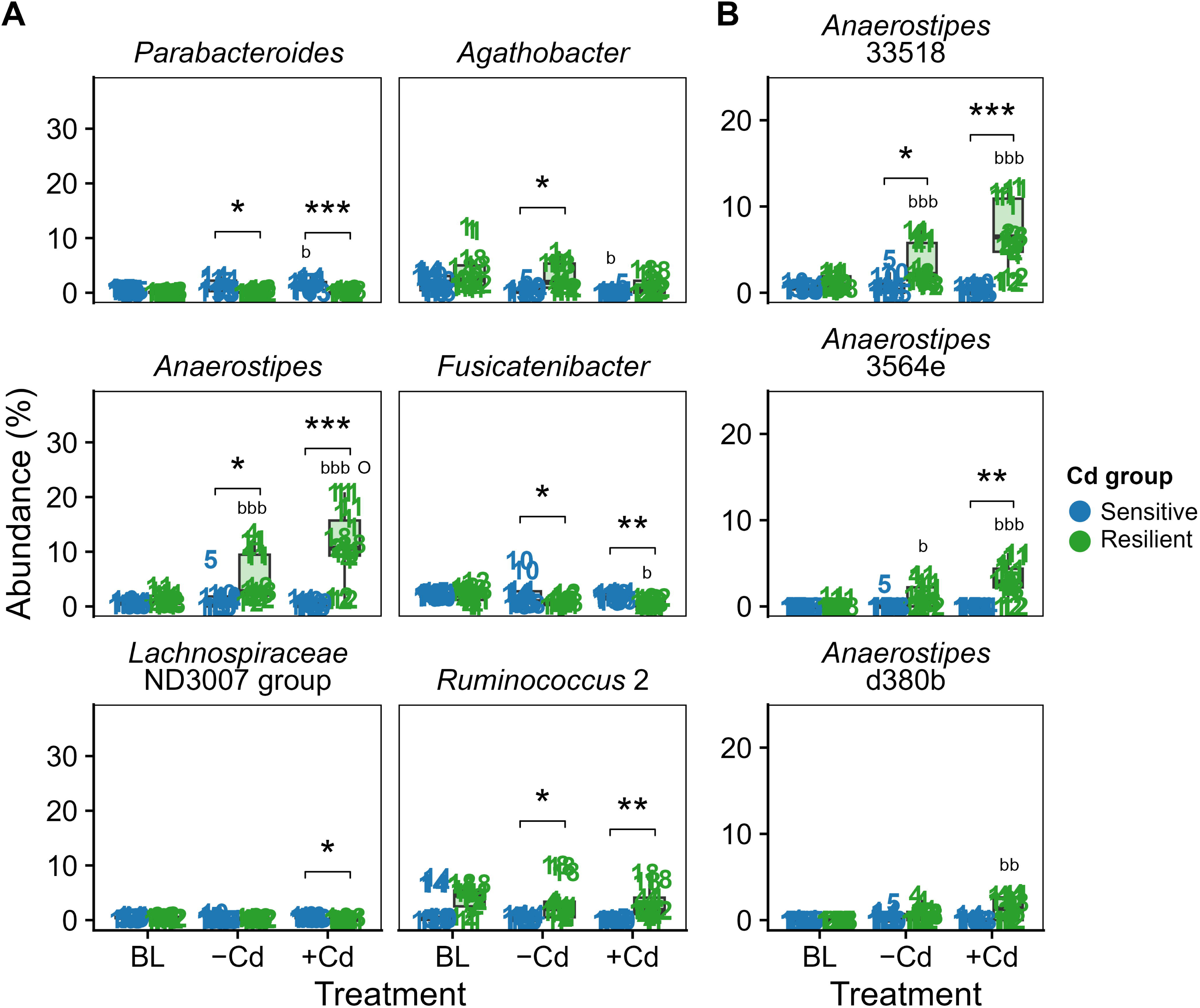
*Anaerostipes* abundance increased in Cd-resilient microbiomes and not in Cd-sensitive microbiomes. A) Abundance of genera with significant differences between Cd-sensitive and Cd-resilient microbiomes across one or more conditions; B) abundance of *Anaerostipes* amplicon sequence variants (ASVs). Asterisks denote significant differences between Cd-resilient and Cd-sensitive communities (*, p<0.05; **, p<0.01; ***, p<0.001). “b” annotations indicate significant differences from baseline (BL) within each group (^b^, p<0.05, ^bb^, p<0.01, ^bbb^, p<0.001); “O” annotations indicate significant differences from no Cd (-Cd) within each group (^O^, p<0.05). Significance of differences were determined with Wilcoxon’s test with Benjamini-Hochberg p-value adjustment implemented after centered log-ratio transformation of raw read counts implemented within the ALDEx2 R-package.

Three *Anaerostipes* ASVs were detected across the communities (Figure 6B). The resilient communities contained all three ASVs, and each exhibited a significant increase in abundance during growth in the presence of Cd. The sensitive communities contained detectable levels of only two of the three ASVs. At baseline, the abundance of these ASVs were not significantly different from the resilient communities; however, during *in vitro* culture, their abundance did not change significantly in the presence or absence of Cd. A BLAST search of the 16S rRNA gene sequences for these three *Anaerostipes* ASVs was unable to provide definitive species identification, as all three sequences had the highest match (>99.5%) to the same two *Anaerostipes* strains (*Anaerostipes amylophilus* H1_26 .13414_6_14.5 and *Anaerostipes hadrus* BA1).

### Anaerostipes abundance was associated with butyrate production in resilient communities during Cd stress

To further understand linkages between changes in microbiota composition and butyrate production, we identified genera that were linearly associated with butyrate production under - Cd and +Cd conditions in resilient and sensitive microbiotas (Figure 7). Butyrate production in the resilient communities strongly correlated with two genera known to include butyrate-producing species: *Faecalibacterium* in the absence of Cd and *Anaerostipes* in the presence of Cd. Butyrate production was also correlated to an unknown genus in the *Lachnospiraceae* family in the presence of Cd; several butyrate-producing species are members of the *Lachnospiraceae* family. In contrast, three genera were associated with butyrate production in the absence of Cd in the sensitive communities: *Coprococcus,* a genus known to contain butyrate-producing species, and two genera that are not known to contain butyrate-producing genera (*Escherichia-Shigella* and *Dorea*). When Cd was present, 13 genera were associated with butyrate production including genera known to include butyrate-producing species (*Blautia*, *Oscillibacter, Ruminoclostridium* 6, *Ruminococcaceae* UCG-002, and *Subdoligranulum*) as well as several other genera not known to include butyrate-producing bacteria. The large number of correlations with taxa not known to produce butyrate likely reflect the higher number of microbial interactions in sensitive communities and their reduced production of butyrate.

**Figure 7.**
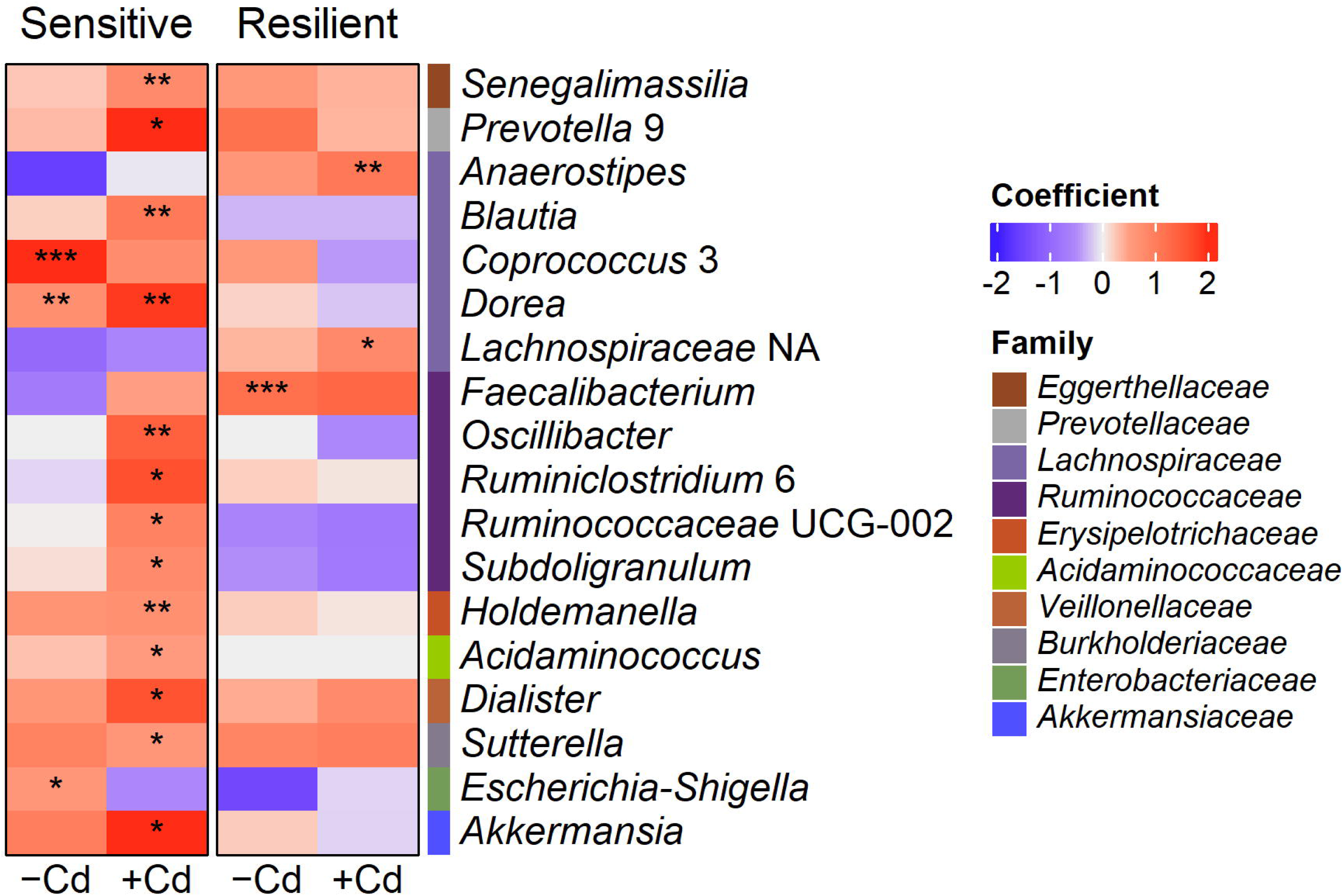
Different genera were associated with butyrate in Cd-sensitive and Cd-resilient microbiomes with (+Cd) or without (-Cd) cadmium stress. Heatmap of the linear associations between butyrate concentrations and genera abundances under +Cd and -Cd conditions by Cd-sensitivity group; the colors represent the effect size and direction of the association; only genera with at least one significant positive association are shown; * q<0.05, ** q<0.01, *** q<0.001; Microbiome Multivariable Association with Linear Models (MaAsLin2).

### Cd-resilient communities had smaller networks of genera that interacted with Anaerostipes that were more conserved under Cd stress

To further investigate potential factors influencing differences in microbiota composition between resilient and sensitive communities, we identified *Anaerostipes* interaction networks based on Spearman correlations between genera in sensitive and resilient communities in the presence and absence of Cd. Network analysis revealed large differences in *Anaerostipes* networks between sensitive and resilient microbiotas (Figure 8). In sensitive communities without Cd stress, the network contained 349 total connections, with nearly all of them positive (324 positive; 25 negative). With Cd stress, only 95 (27%) of these connections were observed, indicating substantial differences in network configurations in the presence and absence of Cd. In contrast, the network in the resilient communities in the absence of Cd contained far fewer connections (60), with 56 positive and 4 negative correlations. In resilient communities cultured with Cd, 24 (40%) of the correlations observed in the absence of Cd were also detected. In the presence of Cd, there were four genera that were directly associated with *Anaerostipes* in resilient communities: [Ruminococcus] gauvreauii group, *Senegalimassilia*, and two genera from the *Erysipelotrichaceae* family—*Catenibacterium* and *Holdemanella*.

**Figure 8.**
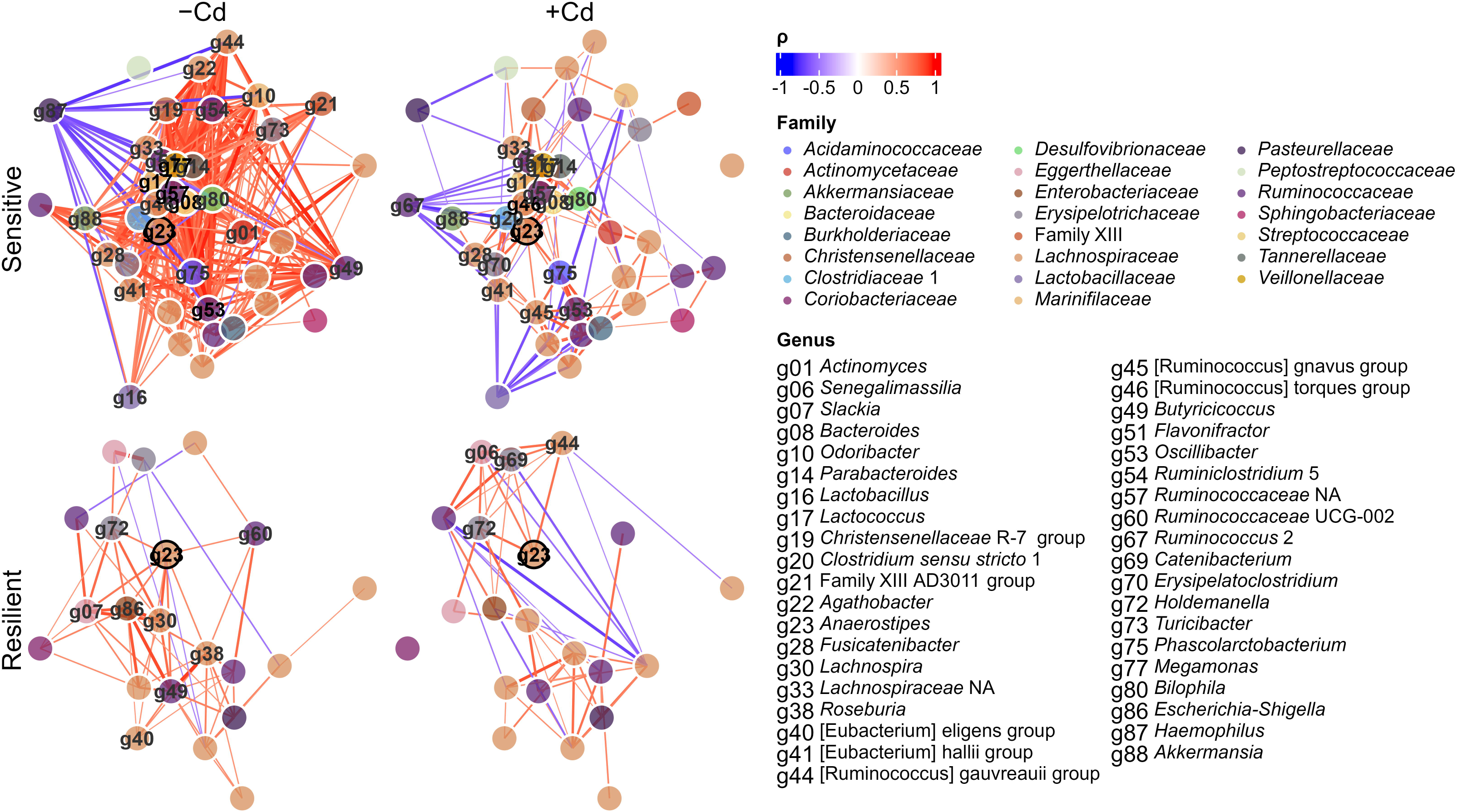
*Anaerostipes* network plots of sensitive and resilient microbiotas with (+Cd) and without (-Cd) cadmium stress. Networks are based on Spearman correlations among genera; only |ρ|>0.5 are plotted. The darkness and color of the lines indicates the effect size and direction of the association; only genera that are directly linked to *Anaerostipes* (g23) are labeled; other taxa are just colored by family.

### Supplementation with Anaerostipes or Cd-resilient communities can restore butyrate production in Cd-sensitive communities

To evaluate the potential of Cd-resilient communities or *Anaerostipes* to restore butyrate production to sensitive microbiotas under Cd stress, two Cd-resilient microbiotas (1 and 12) and two species of *Anaerostipes* (type strains of *A. hadrus* and *A. caccae*) were selected for co-culture experiments with Cd-sensitive microbiotas (13 and 15). The resilient microbiotas were selected based on variable *Anaerostipes* abundance (Figure 9A) and butyrate production (Figure 9B) from our original experiment. A comparison of butyrate production in sensitive microbiotas, with or without supplementation with resilient communities (∼10% total abundance), revealed that both resilient communities increased butyrate production in sensitive microbiota 13, and the magnitude of the increase corresponded to *Anaerostipes* abundance in the resilient community (Figure 9C). However, neither resilient community restored butyrate production to sensitive microbiota 15. Remarkably, a co-culture of either sensitive microbiota with *Anaerostipes* (∼10% total abundance) resulted in the highest butyrate production across all treatments.

**Figure 9.**
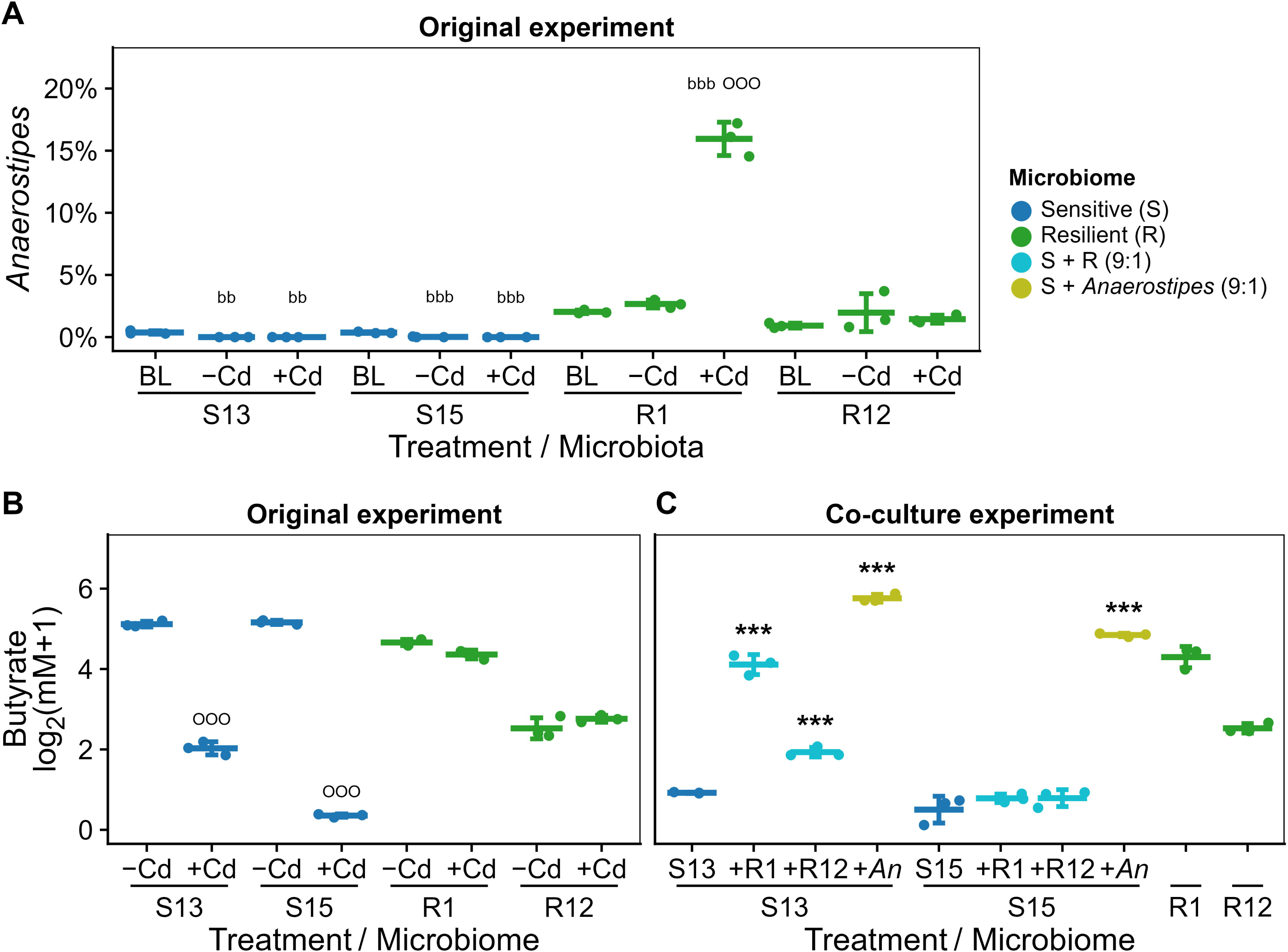
*Anaerostipes* strains increased butyrate production in sensitive microbiotas under Cd exposure. A) Relative abundance of *Anaerostipes* in selected sensitive (S13, S15) and resilient (R1, R12) microbiotas at baseline (BL) and after 24 hours without (-Cd) or with (+Cd) cadmium; data re-plotted form Fig. 6; B) butyrate production of selected sensitive (S13, S15) and resilient microbiotas (R1, R12) without (-Cd) and with (+Cd) cadmium after 24 h of culturing; data replotted from Fig. 3; C) butyrate production of sensitive microbiotas (S13, S15) measured after 24 h with Cd (+Cd) and in co-culture with resilient microbiotas (R1, R12) or *Anaerostipes* species mix (*An*); ^bb^p<0.01, ^bbb^p<0.001 compared with baseline (BL); ^OOO^p<0.001 compared with -Cd; ***p<0.001 compared with sensitive microbiome alone (Tukey’s HSD within microbiome, p < 0.05).

### Cd resilience not correlated with levels of Cd binding by communities

To investigate Cd binding and sequestration by the microbiota as a potential mechanism underlying Cd-resilience, we quantified Cd distribution in cultures following fermentation in the presence of Cd across the full set of microbiota samples. Very little Cd was free in solution, with almost all the Cd tightly or loosely bound to cells (Table 1). However, there were no significant differences in free or bound Cd between resilient and sensitive microbiotas. These results indicate that Cd-binding capacity of the microbiota is likely not linked to the persistence of butyrate production under Cd exposure.

**Table 1.** There were no differences in Cd binding between Cd-resilient and Cd-sensitive microbiomes. Percent of total Cd after 24 h of fermentation with Cd that was free, bound to cells, or loosely adherent in the culture medium.

| Fraction | Sensitive | Resilient | p-value* |
| --- | --- | --- | --- |
| Free | 0.656 ± 0.462 % | 1.45 ± 1.38 % | 0.18 |
| Bound to cells | 38.2 ± 9.68 % | 38.1 ± 11.0 % | 0.69 |
| Loosely adherent (by difference) | 61.1 % | 60.5 % |  |

\*p-value corresponds to Sensitive = Resilient within row, Wilcoxon test.

### Predicted enrichment of stress response pathways in Cd-resilient communities under Cd stress

Given that Cd binding capacity did not explain differences between resilient and sensitive microbiotas, PICRUSt2 was used to further investigate potential alternative mechanisms by predicting differences in microbiota functional potential. Very few differences in predicted pathways between sensitive and resilient communities were evident in fecal samples (Figure 10A). After 24 h culture, significantly more predicted pathways were elevated in the sensitive communities compared with the resilient microbiotas in the absence of Cd (Sensitive = 45 pathways significantly higher versus resilient = 24, χ^2^ = 6.39, p = 0.012). Under Cd stress, the opposite trend was found: significantly more predicted pathways were elevated in the resilient communities compared with the sensitive microbiotas (Sensitive = 9 pathways significantly higher versus resilient = 21, χ^2^ = 4.8, p = 0.029). The predicted pathways that were elevated in the resilient communities under Cd stress suggested enrichment of microbes with stress defense mechanisms, e.g., methionine biosynthesis, menaquinone and demethylmenaquinone biosynthesis, and teichoic acid (poly-glycerol) biosynthesis, while the predicted pathways that were elevated in the sensitive communities suggested a focus on rapid metabolic turnover, e.g., biotin and pyridoxal-5′-phosphate (vitamin B6) biosynthesis/salvage, glycolysis/Entner–Doudoroff, reductive TCA cycle, and fermentative propanoate production, and ADP-L-glycero-β-D-manno-heptose biosynthesis (Figure 10B).

**Figure 10.**
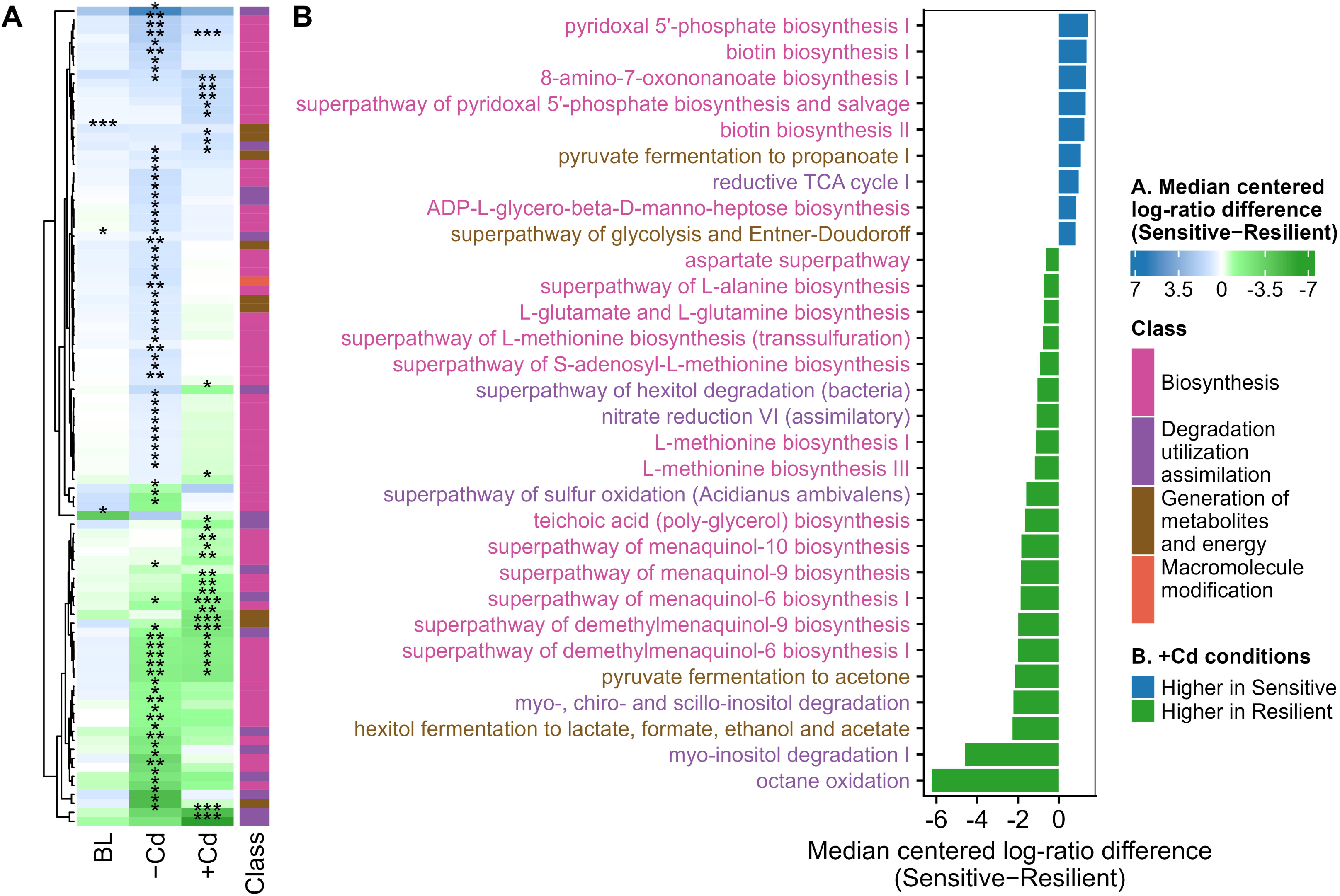
Predicted pathways related to stress defense enriched in Cd-Resilient communities under Cd stress. A) Heatmap showing predicted pathways (PICRUSt2) with significant differences between Cd-sensitive and Cd-resilient microbiotas in fecal samples at baseline (BL) and after 24 h cultures without (-Cd) or with (+Cd) Cd; asterisks denote significant differences between Cd groups (* p<0.05, ** p<0.01, *** p<0.001); heatmap is colored by the Cd group for which they were significantly enriched, with the darkness of the color indicating the magnitude of difference. Statistical significance was determined by Wilcoxon’s test with Benjamini-Hochberg p-value adjustment implemented after centered log-ratio transformation of raw read counts implemented within the ALDEx2 R-package; B) relative abundance of predicted pathways with significant differences under Cd stress; pathway names colored by class of pathway.

## Discussion

In accordance with previous literature (14, 26–29), we observed that Cd stress can be profoundly detrimental to butyrate production by the microbiota using 24-hour human fecal cultures. However, we also identified varying magnitudes of response to Cd stress: some microbiomes exhibited only minor decreases in butyrate production and were classified as Cd-resilient, while others exhibited large decreases in butyrate production and were classified as Cd-sensitive. Analysis of microbiota composition identified a key butyrate-producing genus, *Anaerostipes*, that was significantly enriched in Cd-resilient communities compared to Cd-sensitive communities and whose abundance was highly correlated with butyrate production under Cd stress. Consistent with this linkage between *Anaerostipes* levels and butyrate production by resilient communities under Cd stress, we observed that supplementation of Cd-sensitive microbiotas with *Anaerostipes* could restore butyrate production under Cd stress.

We also observed an inverse relationship between butyrate and lactate levels during Cd stress, with lactate levels significantly higher in sensitive microbiomes cultured in the presence of Cd for 24 h compared to either sensitive communities cultured in the absence of Cd or resilient communities cultured in the presence or absence of Cd. In healthy adults and children, lactate is produced by specific colonic bacteria, but is further metabolized into butyrate (and propionate) such that overall lactate concentrations stay relatively low (30, 31). In adults and children, high lactate concentrations and low gastrointestinal pH is associated with gastrointestinal disorders such as inflammatory bowel disease (32) and lactic acidosis (33). The importance of lactate-utilizing bacteria of the gut microbiome in converting lactate and acetate into butyrate has also been supported by experiments from batch culture and *in silico* models (34–36). In sensitive microbiomes exposed to Cd, the accumulation of lactate likely reflects a disruption in this metabolic conversion due to the loss of key lactate-utilizing taxa. Sensitive microbiomes also had elevated concentrations of acetate after culturing in the presence of Cd which is likely related to the reduction in lactate-utilizing Clostridia, many of which also use acetate in the generation of butyrate (24, 34, 36, 37).

In addition to these significant impacts on secondary fermenters that utilize lactate and some acetate to produce butyrate (38) in Cd-sensitive communities, our data also showed that Cd-treatment impacted primary fermenters responsible for direct utilization of carbohydrates—more so in Cd-sensitive communities than Cd-resilient. Reduced carbohydrate utilization has been suggested in many studies to lead to negative health outcomes to the host (39–42).

Previous studies have shown that Cd exposure often reduces the abundance of beneficial gut bacteria, including *Lachnospiraceae*, *Ruminococcaceae*, and *Prevotellaceae*, which are crucial to maintaining gut barrier function, metabolic balance, and immune modulation and include both primary and secondary fermenting species (14, 26–29). In our study, Cd-resilient communities contained higher abundances of *Lachnospiraceae* in the presence of Cd than the Cd-sensitive communities. Higher levels of *Lachnospiraceae* in resilient communities were primarily due to higher levels of *Anaerostipes* in Cd-resilient compared with Cd-sensitive communities. While *Anaerostipes* levels were not different at baseline between resilient and sensitive groups, levels of *Anaerostipes* increased in resilient communities during cultivation, both in the presence and absence of Cd. This was, in fact, the only genus with significant increases beyond baseline among all *Lachnospiraceae* in any community. *Anaerostipes* contain species that utilize lactate and acetate to produce butyrate (43, 44). The lack of a corresponding increase in *Anaerostipes* in the sensitive communities under Cd stress may partially explain the higher levels of lactate observed in Cd-sensitive communities under Cd stress.

The reason for the enrichment of *Anaerostipes* in the Cd-resilient communities and not the Cd-sensitive communities is not known, as there were no significant differences in levels of *Anaerostipes* between Cd-resilient and Cd-sensitive communities at baseline. Network analysis revealed distinct interaction networks with *Anaerostipes* in Cd-resilient and Cd-sensitive microbiomes; *Anaerostipes are* part of an extensive interconnected network in Cd-sensitive communities and part of a sparse network in Cd-resilient communities. Low-abundance taxa are often inferred to interact with many other taxa, either because these taxa provide a keystone function to the community or because of a technical artefact of the inference method (45). As a majority of microbial interactions in Cd-sensitive communities were lost under Cd stress, the data suggest that *Anaerostipes* was part of a Cd-sensitive network of bacteria that limited the adaptive capacity of *Anaerostipes* to persist in the presence of Cd (46, 47).

The sparse *Anaerostipes* interaction network in Cd-resilient communities suggests less interdependence among taxa that could limit *Anaerostipes* growth. Furthermore, a greater proportion of these connections were maintained in the presence of Cd, suggesting that these microbiotas exhibit enhanced structural resilience and functional stability under Cd stress. Among the limited connections to *Anaerostipes*, two belonged to the *Erysipelotrichaceae* family, *Catenibacterium* and *Holdemanella*, potentially indicating a role for this family in supporting the survival of *Anaerostipes* in the resilient communities under Cd stress. *Erysipelotrichaceae* has been observed at elevated levels—alongside several *Lachnospiraceae* species—in the microbiota of individuals living near smelting mines, where exposure to heavy metals is elevated (48). While these findings suggest that *Anaerostipes* and *Erysipelotrichaceae* interactions may play a role in adapting to Cd stress, further characterization of microbes in this family are needed to better understand the mechanisms behind any protective interaction.

Alternatively, the reason for the persistence of *Anaerostipes* in the Cd-resilient communities and not the Cd-sensitive communities could be due to differences in Cd-tolerance among *Anaerostipes* strains. Our data suggested that the composition of the *Anaerostipes* genus differed between Cd-sensitive and Cd-resilient communities, with three ASVs present in resilient communities and only two in Cd-sensitive. However, 16S rRNA gene sequences for these ASVs were not enough to decipher any functional differences in Cd tolerance among these ASVs. No studies have reported the strain-level variation in Cd or heavy metal tolerance among *Anaerostipes*, although distinct carbohydrate-utilization, amino acid-metabolism, and SCFA production genomic diversity has been shown (49, 50). Importantly, Wang et al. found increased *Anaerostipes* abundances in the microbiotas of ducks after Cd exposure (51), and Zou et al. found higher abundances of *Anaerostipes* in environmental samples from coal mines that were high in heavy metal (lead) compared with mines that were low in heavy metal (52).

Two findings from our study suggest direct tolerance of *Anaerostipes* to Cd, at least for some strains. First, co-culture experiments demonstrated that the addition of *Anaerostipes* strains to sensitive microbiomes under Cd stress was able to dramatically enhance butyrate production in these communities. Second, in resilient communities, butyrate production was primarily associated with *Faecalibacterium* in the absence of Cd and *Anaerostipes* in the presence of Cd. This suggests that butyrate production may have shifted from *Faecalibacterium* to Cd-tolerant *Anaerostipes* in Cd-resilient microbiomes.

However, there are unexplained factors that govern the relationship between *Anaerostipes* abundance and butyrate production under Cd stress. For example, the addition of resilient microbiomes containing varying levels of *Anaerostipes* to sensitive communities under Cd stress did not consistently rescue butyrate production. One possible explanation is that ecological competition between certain sensitive and resilient communities may lead to incompatible community assembly or competitive exclusion within shared ecological niches, thereby limiting the establishment of Cd-tolerant *Anaerostipes* with the accompanying restoration of butyrate production (53).

Numerous bacterial heavy metal stress response strategies have been described, including extracellular barriers, intracellular and extracellular sequestration, active efflux of metal ions, enzymatic reduction, and biomethylation (54). Initially, we hypothesized that Cd tolerance might be related to the ability of some members of the community to sequester Cd, making these ions unavailable to exert harmful effects on sensitive members of the gut microbiota (54–57). This, in turn, would allow sustained production of butyrate. However, we found that Cd resilience, as indicated by butyrate production, was not directly linked to Cd binding capacity.

Instead, predicted functional analysis suggested that resilient microbiotas were enriched in taxa encoding stress responses pathways that could provide protection against Cd stress, whereas Cd-sensitive microbiotas were enriched in pathways more susceptible to Cd stress. Higher predicted abundance of methionine biosynthesis in Cd-resilient communities could reflect an enhanced capacity for sulfur assimilation and thiol/redox homeostasis (58) that could limit oxidative stress caused by Cd-mediated disruption of thiol pools (59). Further, higher abundance of menaquinone and demethylmenaquinone biosynthesis pathways in Cd-resilient communities could indicate a greater capacity to sustain respiratory electron transport and redox balance under metal stress (60) by stabilizing electron flow through the membrane and compensate for Cd-sensitive respiratory components. In contrast, Cd-sensitive microbiomes were enriched for pathways associated with biosynthesis and rapid metabolic turnover, such as biotin and pyridoxal-5′-phosphate (vitamin B6) biosynthesis/salvage and central carbon pathways. Cd is known to disrupt redox- and metal-sensitive enzymes (including Fe–S proteins), which may render highly active metabolic networks more vulnerable to Cd-induced dysfunction (61). Consistent with this interpretation, Cd-sensitive communities also showed higher predicted abundance of lipid and envelope-associated pathways. Because Cd can damage membrane-associated systems and interfere with metalloproteins, enrichment in these pathways may further increase susceptibility to metal stress (62).

Although we observed distinct Cd phenotypes among microbiomes, ranging from Cd-sensitive to Cd-resilient communities, our study is not without limitations. First, the molecular and physiological bases for this distinction remain unclear, likely reflecting substantial variability among communities. Notably, resilience does not appear to be driven by a single mechanism, as different resilient microbiomes may rely on distinct strategies to cope with Cd stress (54). Our study suggested that direct sequestration of Cd by the gut microbiota is likely not related to Cd-resilience but instead is related to having structurally resilient microbial networks or presence of Cd-tolerant butyrate-producing taxa. It is plausible that one, both, or additional, unidentified mechanisms, are employed by members of the resilient microbiomes to prevent Cd toxicity. Future investigations employing metagenomic, transcriptomic, or culture-based approaches—likely involving defined communities or specific strains of bacteria, from *Anaerostipes*, for example, will be essential to elucidate the specific pathways involved in Cd resilience.

Our study also did not quantify Cd in fecal samples or attempt to quantify the environmental exposure of fecal donors to Cd. These data would be valuable for determining whether prior Cd exposure shapes the functional response of the gut microbiome to Cd stress. It is plausible that microbiomes that are exposed to higher levels of Cd develop a tolerance to Cd. The observed variability under controlled *in vitro* conditions suggests that intrinsic microbial community structure and metabolic potential may play a role independent of baseline Cd levels. Integrating Cd exposure metrics into fecal donor metadata in future work will be critical to understanding these effects.

Our study also relied on a single Cd concentration and a single exposure time to differentiate between sensitive and tolerant communities. In our previous, continuous-culture study, we did not find crossover effects between the Cd-sensitive and Cd-resilient communities over time (25). However, we did find differences in Cd-tolerance among bacterial strains isolated from a Cd-resilient microbiota when the Cd concentration was varied. In this study, we selected a physiologically relevant Cd concentration of 20 (mg/L) to test the effects of Cd on the gut microbiota (63, 64). However, based on our calculations, physiologically relevant concentrations could range from 1.36-51.5 mg/L. Therefore, we acknowledge that further studies incorporating more Cd concentrations would provide insight into the variability in response of microbial communities to Cd stress.

Despite these limitations, our findings clearly demonstrate the existence of both Cd-resilient and Cd-sensitive microbiomes. Our study points to lactate-utilizing, butyrate-producing bacteria as important for the separation of Cd-sensitivity and -resilience, with *Anaerostipes* playing an important, albeit not completely defined, role. By identifying compositional features that distinguish sensitive from resilient microbiomes, this work deepens our understanding of how gut microbial interactions may counteract the adverse effects of Cd exposure. Together, these findings provide a foundation for future mechanistic studies aimed at protecting the gut microbiota from Cd toxicity.

## Materials and methods

### Fecal sample collection and preparation

Fecal samples were collected from twenty healthy adults with no history of gastrointestinal disorders or probiotic/antibiotic consumption within the last six months. The donors ages ranged from 23-58 years old (31.0 ± 10.8) and there were 7 males and 13 females. The study used fecal samples as a source of gut bacteria; it did not include any dietary intervention for the participants. The Institutional Review Board of the University of Nebraska-Lincoln approved all procedures involving human subjects prior to the study (approval number 20210621206EP). All subjects provided written informed consent to participate. Originally, 21 fecal samples were collected and numbered from 1-21. Subsequently, it was discovered that the fecal donor of microbiotas 2 and 7 were the same person, so microbiota 7 was removed from the data analysis.

Fresh fecal samples were mixed with anaerobic sterile phosphate-buffered saline (pH 7.0) containing 10% glycerol at a ratio of 1:9 w/v in sterile filter bags (Filtra-Bag, Labplas, Sainte-Julie, QC, Canada) and mixed with a Stomacher LabBlender 400 (Seward, London UK) for 3 minutes. The resulting slurry was transferred to an anaerobic chamber (Bactron X, Sheldon Manufacturing, Cornelius, Oregon, USA with 5% H_2_, 5% CO_2_, and 90% N_2_ atmosphere), aliquoted into 15 mL tubes, and stored at -80 °C until use.

### -Cd media preparation

Fermentation medium that mimics the nutritional environment of the colon (65) was used to culture fecal microbiomes. To prepare the fermentation medium without Cd (-Cd), 0.8 g each of arabinogalactan (from Larch wood, A1328, TCI), cellobiose (Acros Organics), inulin (Beneo Orafti), soluble starch (Acros Organics), pectin (J6102, Alfa Aesar), and xylan (from Corn core, X0078, TCI); 2.0 g of yeast extract (BP9727, Fisher Scientific); 2.0 g of peptone (M-18664, Fisher Scientific); 0.5 g of bile salts (LP0055, OXOID); 0.01 g of magnesium sulfate; 0.01 g of calcium chloride; 3.0 g of sodium bicarbonate; and 2 mL of Tween 80 were mixed in distilled water. The pH was adjusted to 6.8 using 6M hydrochloric acid, and the solution volume was adjusted to 900 mL with distilled water. The mixture was autoclaved at 121°C for 30 minutes. In parallel, a second solution was prepared by adding 0.5 g of cysteine hydrochloride monohydrate, 1.920 µL of acetic acid (A6283, Sigma), 95 µL of propionic acid (60-047-009, Fisher), 700 µL of butyric acid (AC108111000, Fisher), 93 µL of isobutyric acid (AAL04038AE, Fisher), and 109 µL of isovaleric acid (AC156691000, Fisher) to 50 mL of distilled water. The pH was adjusted to 6.8 with 10M sodium hydroxide, and the volume was adjusted to 98 mL with distilled water. This solution was filter sterilized and added aseptically to the autoclaved medium. Additionally, 1 mL of a vitamin K solution (5 mg/mL in ethanol) and 1 mL of a hemin solution (0.5% w/v in DMSO) were filter-sterilized and added to the medium, resulting in a final volume of 1 L. The media were placed in an anaerobic chamber and pre-reduced for 72 hours before fermentation.

### +Cd media preparation

For the Cd-containing medium (+Cd), 32.6 mg of CdCl_2_ was added to fermentation medium per liter achieve a final concentration of 20 mg/L Cd. Cd concentrations in human fecal samples have been reported to range from 0.125 mg/g to 6.33 mg/g (wet weight) (63, 64). Considering an average daily fecal output of approximately 122 g per person wet basis (66), this corresponds to a total daily Cd excretion of 15.3 mg to 772 mg. To simulate physiologically relevant exposure levels *in vitro*, these values were normalized to reflect estimated colonic Cd concentrations, resulting in a working range of approximately 1.36 to 51.5 mg/L based on a colonic volume that ranges from 0.6 to 3.0 liters (67). A median concentration of 20 mg/L was selected as it falls within this estimated physiological range and represents a plausible exposure level for populations living in regions with elevated environmental Cd contamination. In addition, this level was chosen to approximate a transient acute luminal exposure, enabling assessment of how distinct microbial communities respond to a biologically meaningful Cd challenge. This concentration is also supported by previous studies showing significant microbiota shifts at 20 mg/L Cd (25) and aligns with reported dietary and fecal Cd levels (63).

### In vitro cultures

*In vitro* cultures were performed using fecal slurries from 20 donors as sources of gut microbiota. Culturing was performed in triplicate, in media with (+Cd) and without (-Cd) Cd. Tubes with 6.0 mL of pre-reduced fermentation media were inoculated with 0.6 mL of fecal slurry, capped, and incubated at 37°C with orbital shaking (125 rpm) for 24 h. Samples were collected after 0 h (baseline) and 24 h (-Cd and +Cd) of fermentation and immediately stored at - 80°C.

### Microbial metabolite analysis

SCFA were analyzed by gas chromatography as described by (68). Briefly, fermentation slurries were centrifuged 10,000*g* for 5 min. The supernatant (0.4 mL) was then mixed with 0.1 mL of 7 mM 2-ethylbutyric acid (109959, Sigma) in 7 M potassium hydroxide, 0.2 mL of 9 M sulfuric acid, and approximately 0.1 g of sodium chloride in a 2 mL tube. SCFA were extracted by adding 0.5 mL diethyl ether, vortex mixing, and recovering the top layer into a glass vial. Four microliters of this diethyl ether extract were injected into a gas chromatograph (Clarus 580; PerkinElmer, MA, USA) equipped with a capillary column (Nukol, 30 m length x 0.5 mm inner diameter, 0.25 μm film thickness, Supelco, PA, USA) and detected with a flame ionization detector.

Quantification of lactic acid was performed using supernatants from *in vitro* culture samples as described for SCFA, except the supernatants were 1:10 diluted prior to analysis. Diluted supernatants were filtered through a 0.45 μm nylon filter and then injected into an HPLC system (Agilent 1260 Infinity, Waldbronn, Germany) equipped with an Agilent 1260 Infinity quaternary pump (G1311B), an autosampler (G1367E) and a diode array detector (G4212B) set at a wavelength of 210 nm, using 360 nm as reference. Separation of lactic acid was performed using an Aminex HPX-87H Organic Acid Analysis Column (300 x 7.8 mm, Bio-Rad) with 10 mM sulfuric acid as the mobile phase. The column temperature was set to 50 °C, with isocratic elution at a flow rate of 0.6 mL/min and a run time of 35 min. Lactic acid was quantified as the sum of D- and L-lactic acid using standard calibration curves prepared from authentic standard solutions in the mobile phase.

To calculate metabolite production after 24 h, we subtracted the initial metabolite concentration (0 h, or baseline) from the final concentration (24 h). This resulted in some negative values, indicating that metabolite utilization exceeded production. To determine the log_2_-fold change between +Cd and -Cd, we added a constant (one plus the absolute value of the lowest number) to all data points to eliminate negative numbers and zeros before calculating the log_2_-fold change on the transformed values.

### Carbohydrate degradation

Carbohydrate degradation was calculated as the difference in carbohydrate concentration between 0 and 24 h of fermentation. Carbohydrate concentration was measured from the supernatant of sampled *in vitro* fermentations (centrifuged at 10,000*g* for 5 minutes). Arabinogalactan, cellobiose, soluble starch, and pectin were quantified from their constituent sugars after acid hydrolysis using HPLC. Briefly, supernatant (0.25 mL) from each sample was mixed with 0.5 mL of internal standard (inositol, 1 mg/mL), 0.1 mL of 12 M sulfuric acid, and 2.15 mL of water, and then autoclaved at 15 psi for 1 h. For neutral sugar analysis, the mixture was then neutralized with calcium carbonate until bubbling stopped, then passed through a 0.45 µm filter for HPLC injection. For uronic acid analysis, the samples were filtered through a 0.45 µm filter without neutralization.

Sugar quantification was performed using the HPLC system described previously, except a refractive index detector (G1362A) was used. Separation of neutral sugars was achieved with an Aminex HPX-87P Column (300 x 7.8 mm, Bio-Rad) using water, which was autoclaved and filtered with a 0.45 µm Nylon filter, as the mobile phase. The column temperature was maintained at 80°C, with isocratic elution at a flow rate of 0.6 mL per minute and a sample injection time of 35 minutes. Sugars were determined using external standard calibration curves prepared from solutions of glucose, arabinose, galactose, and xylose relative to the internal standard. Separation of uronic acids was performed as was previously described for lactic acid. Galacturonic acid was quantified using external standard calibration curves prepared from standard solutions of galacturonic acid relative to the internal standard.

Fructose from inulin was not well quantified from the above chromatographic analysis, likely because of the harsh hydrolysis conditions. Therefore, inulin was quantified separately using a Megazyme Fructan Assay Kit (Fructan Assay Kit, K-FRUC; Megazyme Ltd., Bray, County Wicklow, Ireland) following the manufacturer’s instructions.

### Microbiota composition

Aliquots (0.2 mL) from fermentation cultures at 0 h and 24 h were collected and centrifuged at 10,000 X g for 5 minutes to recover bacterial pellets. DNA from the fecal samples of all microbiomes (pellets) was extracted using the BioSprint 96 One-For-All Vet kit (Qiagen, Germantown, MD) according to the manufacturer’s instructions. The V4 region of the bacterial 16S rRNA gene was amplified from each sample using the dual-indexing sequencing strategy (69) using the Illumina MiSeq platform and the MiSeq reagent kit v2 (2 × 250 bp) (Illumina, San Diego, CA) according to manufacturer’s protocols in the Nebraska Food for Health Center laboratories.

For 16S rRNA gene sequencing processing, sequences were truncated (220 bases for forward reads and 160 bases for reverse reads) and demultiplexed, and barcodes were removed before sequence analysis with QIIME 2 (70, 71). Quality control, trimming, chimera removal, and denoising were conducted with DADA2 (72), which dereplicated sequences into 100% amplicon sequence variants (ASVs) for exact sequence matching. Taxonomy was assigned using the SILVA 132 database (https://data.qiime2.org/2019.4/common/silva-132-99-515-806-nb-classifier.qza) (73). Samples were rarefied to a sequencing depth of 10,225 reads per sample before analysis. Rarefaction and diversity calculations were performed using the phyloseq package (version 1.48.0) in R (version 4.2.3) (74). For microbiota composition analysis, amplicon sequence variants (ASVs) were filtered to retain only those present in at least 10% of the samples.

### Co-cultures of sensitive gut microbiotas with resilient gut microbiotas or Anaerostipes strains under Cd exposure

To determine if butyrate production could be increased in Cd-sensitive communities under Cd stress, co-cultures with Cd-resilient communities or *Anaerostipes* strains were prepared. Two Cd-resilient microbiotas (1 and 12) and two representative strains of *Anaerostipes*—*A. caccae* (DSM 14662, NCIMB 13811, strain L1-92) and *A. hadrus* (DSM 3319, NCBI 649757, strain VP 82-52)—were selected for co-culture experiments with Cd-sensitive microbiotas (13 and 15).

*Anaerostipes* strains were grown anaerobically at 37 °C for 16 h in pre-reduced brain heart infusion medium supplemented with yeast extract (BHIs; BD 211059, VWR 90000-060; BP9727, Fisher Scientific). Subsequently, individual colonies were inoculated into BHIs broth and incubated overnight under anaerobic conditions. The optical density at 600 nm (OD_600_) of the overnight cultures was measured. Cultures were then serially diluted and plated on BHIs agar to estimate colony-forming units per milliliter (CFU/mL) after 24 h. Based on OD_600_ measurements, cell concentrations for each isolate were estimated.

*In vitro* cultures were performed as described above using Cd-containing medium (20 mg/L), except a 5 mL fermentation volume was used together with a total volume of 0.5 mL of inoculum. Where sensitive and resilient microbiotas were co-cultured, 0.45 mL of the sensitive inoculum and 0.05 mL of the resilient inoculum were used. For treatments including *Anaerostipes*, OD_600_ measurements of overnight cultures were used to estimate cell concentrations. Cultures were centrifuged to obtain cell pellets, which were resuspended in phosphate-buffered saline (PBS). The suspensions were adjusted to 5 × 10 CFU/mL for each strain, and equal volumes of *A. caccae* and *A. hadrus* were combined to prepare the *Anaerostipes* inoculum. Then, 0.45 mL of the sensitive inoculum and 0.05 mL of the *Anaerostipes* mixture were used.

### Cd quantification

Cd concentrations were quantified in both the culture supernatant and bacterial pellet after 24 h of fermentation in Cd-containing medium. For supernatant measurements, samples were centrifuged at 10,000×*g* for 5 min, and Cd levels were measured in the digested supernatant from the 20 microbiomes. For analysis of Cd in the bacterial pellet, bacterial pellets recovered from DMSO stocks were washed three times with PBS buffer containing 1 mM EDTA, followed by centrifugation (10,000×*g* for 1 min) and removal of the rinse solution to remove soluble and loosely adherent Cd to the bacterial cells before quantification. The pellets were digested with 100 µL of nitric acid at room temperature, with 2 hours of incubation at 60°C.

Quantification was performed using inductively coupled plasma mass spectrometry (ICP-MS). Prior to analysis, the supernatants and digested pellets were diluted 10-fold with 2.0% nitric acid, and gallium was added as an internal standard at a final concentration of 50 ppb. Sixty microliters of the diluted supernatant were injected into an ICP-MS instrument (7500cx ICP-MS, Agilent Technologies, CA, USA) operating in reaction and kinetic energy discrimination mode with helium gas at a flow rate of 1.5 mL/min and hydrogen at 3.5 mL/min. Nitric acid (2.0%) was used for rinsing the system between samples. The Cd concentrations were calculated from the ratios of ^111^Cd to ^71^Ga against an external calibration curve (0-100 ppb).

Loosely adhered Cd was calculated as the difference between the initial Cd concentration (20 mg/L) and the sum of Cd measured in the supernatant and in the bacterial pellet.

### Functional prediction of the microbiota

Functional composition of the gut microbiota was calculated from 16S rRNA gene sequencing data using Phylogenetic Investigation of Communities by Reconstruction of Unobserved States 2 (PICRUSt2) (75–80). Pathway data were categorized using classifications designated in the MetaCyc database (https://metacyc.org/).

### Data analysis

All statistical analyses were performed in R (version 4.2.3) using RStudio (version 2024.4.2.764). For metabolite data, differences between treatment (-Cd versus +Cd) and between Cd group (sensitive versus resilient) were determined using Wilcoxon tests with p-values adjusted using Holm’s procedure. Correlations were calculated using Pearson’s method. For microbiota composition, differences in the β-diversity based on UniFrac distances across Cd groups were compared using PERMANOVA with the adonis2 function in vegan (81). To explore differences in microbial taxonomic composition, taxon level abundances were calculated across all hierarchical ranks using the metacoder package (82). Differential taxon abundances and pathway abundances were then compared using Wilcoxon’s test with Benjamini-Hochberg p-value adjustment after centered log-ratio transformation of raw read counts using the ALDEx2 package (83). Statistical significance was determined as q < 0.05.

To investigate how Cd exposure alters microbial association patterns, Spearman correlation was calculated among genus-level relative abundances under -Cd and +Cd conditions for sensitive and resilient microbiotas. *Anaerostipes* networks were generated using the NetCoMi package (84). Networks were sparsified using a correlation threshold of |ρ| ≥ 0.5 to reduce spurious associations and retain moderate-to-strong relationships. No imputation or additional normalization was applied to the correlation matrices.

To assess the associations between butyrate concentrations and genus-level microbial abundance, multivariable linear modeling was performed using the MaAsLin2 package (85) associations with q < 0.05 were considered statistically significant.

In the co-culture experiment, data were analyzed by microbiome using a one-factor ANOVA where treatment was the factor. To identify significant differences among treatments, Tukey’s Honestly Significant Difference test was used.

Graphs were generated using the ggsignif (version 0.6.4), ggpubr (version 0.6.0), ggh4x (version 0.3.0), metacoder (version 0.3.8), cowplot (version 1.1.3), and ComplexHeatmap packages (82, 86–90). Source data used for figure generation can be found in Supplementary File 1. Taxon abundances of Cd-sensitive and Cd-resilient microbiotas at baseline and without Cd (Cd0) or with Cd (Cd20) can be found in Supplementary Table 1.

## Acknowledgements

We thank Dr. Limei Zhang (University of Nebraska–Lincoln) for her insightful suggestion to perform cadmium quantification in the bacterial pellets from our *in vitro* cultures, which strengthened this work. We thank Dr. Anna Seekatz and Megan Seesengood (Clemson University) for helpful discussions about *Anaerostipes*.

## Author contributions

Conceptualization (C.P., D.R., J.A.); data curation (C.P.); formal analysis (C.P., D.R.); funding acquisition (D.R., J.A.); investigation (C.P., S.L., J.S.); methodology (C.P., J.S., D.R., J.A.); project administration (D.R., J.A.); resources (J.S., D.R., J.A.); supervision (D.R., J.A.); visualization (C.P., D.R.); writing—original draft (C.P.); writing—review & editing (C.P., S.L., J.S., D.R., J.A.). All authors read and approved the final version of the manuscript.

## Ethics statement

The studies involving humans were approved by the University of Nebraska-Lincoln Institutional Review Board (approval number 20210621206EP). The studies were conducted in accordance with local legislation and institutional requirements. The participants provided their written informed consent to participate in this study.

## Data availability

Raw sequence reads from 16S rRNA gene sequencing are available in the Sequence Read Archive under accession number <u>PRJNA1435248</u>. The processed datasets and R code generated during the current study are available in the Zenodo repository at DOI: 10.5281/zenodo.18944206 (DOI will be made public upon acceptance).

## Supplemental material

**Supplementary File:** Source data for Figures 1-10 and Table 1.

**Supplementary Table 1:** Taxon abundances of Cd-sensitive and Cd-resilient microbiotas at baseline and without Cd (Cd0) or with Cd (Cd20)

